# Topological data analysis captures complex behavioral dynamics during naturalistic social interaction between domestic ferrets

**DOI:** 10.64898/2026.07.01.735818

**Authors:** Jared Reiling, Nancy Padilla-Coreano, Dhruvi Patel, Flavio Frohlich, Mengsen Zhang

## Abstract

Capturing naturalistic behavioral dynamics is essential for understanding social interaction in ecologically valid settings. Existing investigations of naturalistic social interaction rely on time-aggregated analysis methods better suited for task-based experiments, which lose the complex, moment-to-moment dynamics exhibited in naturalistic settings. The emerging field of topological data analysis (TDA) provides new tools to characterize fine-grained dynamics in time-series data that cannot be captured by time-averaged methods. The present work utilizes Temporal Mapper, a recently developed TDA specifically tailored to analyzing dynamical systems. Temporal Mapper characterizes complex temporal dynamics as transition networks, where nodes are stable states and edges are transitions between states. Originally designed for human neural time series analysis, here we demonstrate the utility of Temporal Mapper to capture rich animal postural dynamics during naturalistic social interaction. We utilized an existing dataset with 12 video recording sessions of two domestic ferrets (*Mustela putorius furo*) during naturalistic interaction and tracked the postures of animals during social interaction. Ferrets were chosen due to their strong social-cognitive skills and rich postural dynamics for investigating social behavior via posture estimation. Temporal Mapper was then used to represent the postural dynamics as transition networks for each recording session. Here, we found that posture states are significantly smaller and more widespread during active social interaction compared to non-social activities. Additionally, the number of sequential postural states before transitioning to new behaviors is more consistent during active social interaction than non-social activities. Together, our findings suggest that social activity has a broad range of unstable postural states arranged in consistent sequences. Our method, Temporal Mapper, allows for network structure analysis of complex naturalistic data, applicable for characterizing rich dynamics in different species, scales, and paradigms.

## INTRODUCTION

Social interaction is highly complex and dynamic process^1^. It has been argued that mental health and psychiatric disorders should be conceptualized as disorders of social interaction^2,3^. Thus, understanding social interaction lays the foundation for creating new interventions for psychiatric disorders. Prior research primarily examined social behavior using task-based paradigms, such as the three-chamber assay^4^, modified chamber assay^5^, operant task^6^, and sociobox^7^, which involve at least one animal stationary behind a barrier while the subject animal moves freely around the enclosure. These tasks were designed to measure behavioral preference for social novelty and recognition, where the total time spent exploring the stimulus mice is an indicator of preference^4–7^. Additional paradigms with freely interacting animals, such as tone-reward training^8^, are designed for competition settings. Behavioral analysis within these task frameworks was primarily accomplished though measuring the average social time between animals scored by humans or average number of wins during social competition across trials. Even within more naturalistic experimental paradigms, including the open test box exploration^9^, the data analysis focus is on explicitly quantifying average time spent socially interacting across trials. These existing studies analyzed behavioral data via time-aggregated measures where the complex, moment-to-moment changes within social behavior are largely lost. To capture more fine-grained temporal information about social dynamics, new behavioral tracking tools emerged, including DeepLabCut^10–12^, SimBA^13^, and Anipose^14^. These new data analysis tools are designed for animal pose estimation using markerless motion tracking of multiple animals in 2D^10,12,13^ and 3D^11,14^ settings from video recordings. Pose estimation tools create high-dimensional time series data of body segment movement of single or multiple animals. The objective of these fine-grained behavioral tracking methods is for supervised behavioral classification^13,15–26^ which alleviates time-intensive manual human annotation. Nevertheless, the resulting rich behavioral time series were often analyzed using time-aggregated statistics, without capturing the temporal dynamics of behavior during interaction. To capture more temporal information within social interaction, modeling tools such as the Hidden Markov models (HMMs)^27–32^ have been applied to describe behaviors and transitions between behaviors from animal movement time series data. However, HMMs often have a limited number of states^16,33^, such that the resulting transition network is not rich enough for topological analysis. Moreover, the interpretation of HMMs states is not always straight forward in a theoretical sense – for example, it has not been shown that HMM states and transitions correspond to known stable states and state transitions in mathematical models of dynamical systems^34^.

Therefore, new methods are needed that provide a more flexible, dynamical state-aware approach to capture and connect behavioral states during naturalistic social interaction. Topological Data Analysis (TDA) provides a new data-driven approach for modeling fine-grained behavioral dynamics. TDA is a growing field in applied mathematics for developing new tools to capture complex, intricate structures within high-dimensional data^35–37^ and is increasingly utilized in neuroscience^34,38–41^. For example, a specific TDA method Mapper^35,42^ has been applied to characterizing brain dynamics as topological representations (i.e., graphs, and high-dimensional counterparts, known as simplicial complexes) from neuroimaging data^34,38,39^. Mapper provides a simplification and visualization of high-dimensional data in the form of simplicial complexes, where nodes correspond to clusters in the data and edges connect regions with overlapping clusters^42^. However, simplicial complexes do not include temporal information within the construction, leaving the sequence of state transitions unknown. To address this gap, a new TDA tool Temporal Mapper^34^ was created, synergizing the principles of Mapper and dynamical systems theory. In practice, Temporal Mapper provides a data-driven approach for determining the states and transitions and connecting them in a global network. This new method was shown to reconstruct ground-truth transition networks from large-scale nonlinear dynamical systems models of the brain, providing a link between mechanistic modeling and data analysis^34^. Empirically, Temporal Mapper has been used to characterize brain dynamics from human fMRI data, revealing rich relations between brain states within and between tasks^34^. Subsequently, Temporal Mapper has been applied to behavioral time series, revealing fine-grained social-behavioral dynamics during psychotherapy that eluded time-averaged methods^43^.

However, in this work, the behavioral time series was obtained through time-intensive human ratings of patients’ and therapists’ warmth and dominance levels, which are not generalizable to the behavioral analysis of animal interactions. The applicability of Temporal Mapper to animal movement data obtained from pose estimation as a means of characterizing social interaction is unknown.

In the present work, we demonstrate the applicability of using Temporal Mapper to capture rich behavioral dynamics from motion tracking data between freely interacting domestic ferrets (*Mustela putorius furo*). We chose ferrets as a model due to their strong social-cognitive skills^44^ and their ability to communicate through distinct postures and body movements^45^, making them excellent for tracking postures via computer vision from video recording. Here, we leverage Temporal Mapper to represent posture dynamics as transition networks, where nodes are unique posture states and edges are transitions between postures. Our computational framework provides a data-driven solution for fine-grained characterization of complex, high-dimensional behavioral dynamics within naturalistic settings.

## RESULTS

### Posture Time Series Data to Transition Networks

First, we demonstrate how posture tracking can be represented as transition networks using Temporal Mapper. Posture dynamics were captured via computer vision from video recording in which individual body segments were tracked during social interaction. These posture dynamics are high-dimensional time series data, where the location of 8 body segments per animal were tracked. Over time, animals exhibit various types of postural states. For illustrative purposes, an example postural time series and states is shown in Figure 1A. Using only posture time series, Temporal Mapper represented postural states as a transition network where the nodes are unique posture states, and the edges represent the transitions between states. For illustrative purposes, an example transition network is shown in Figure 1B. The nodes are colored based on the behavior occurring within each postural state.

**Figure 1:**
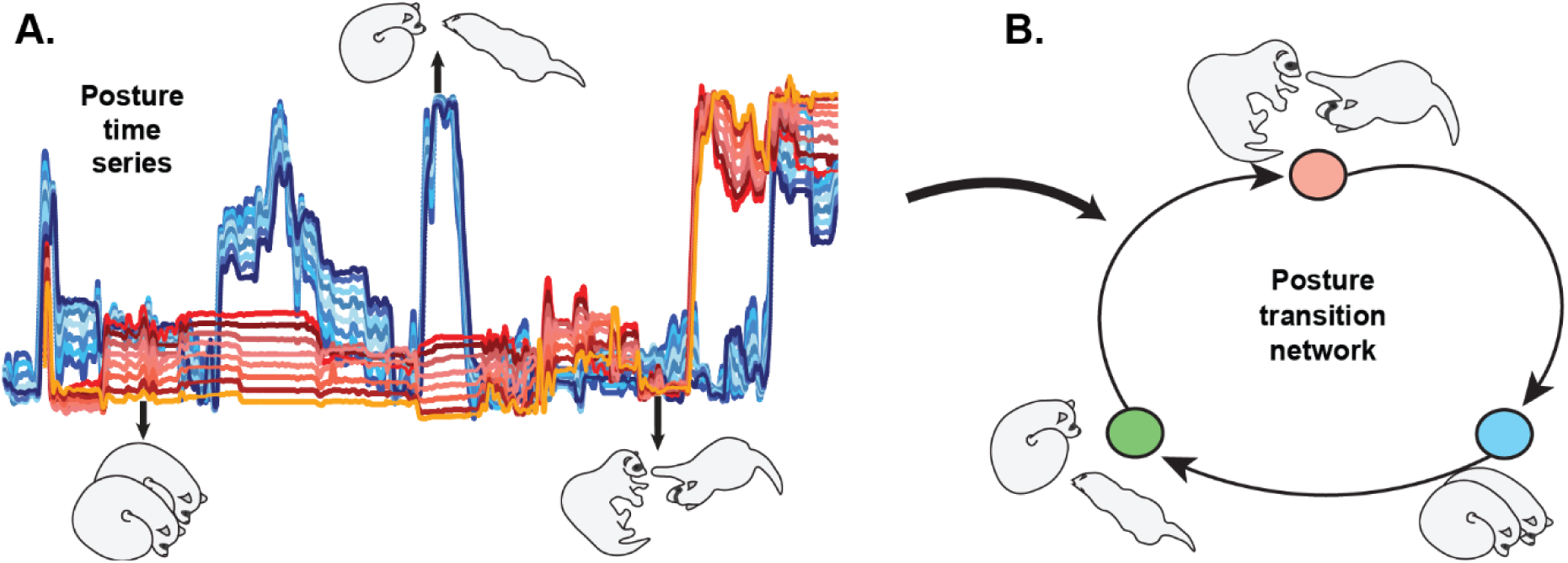
Temporal Mapper represents posture time series as transition network. (A) During naturalistic behavioral recording, animals freely move around the behavioral chamber and display various types of postures over time. In this example, the position of body segments of animal 1 (colored in blue hues) and animal 2 (colored red hues) changes while animals are interacting with each other. For illustrative purposes only, three different postures are illustrated during naturalistic recordings. (B) Temporal Mapper represents postural states as a transition network where nodes are unique postural states and edges are the transitions between postural states. Nodes are colored based on the behavior exhibited during the postural state.

In the present work, we use Temporal Mapper to capture the moment-to-moment postural dynamics during naturalistic social interaction. The following sections demonstrate the utility of Temporal Mapper to characterize social interaction between freely interacting ferrets.

### From Conceptual Framework to Application: Acquiring Relative Posture Time Series Data via Computer Vision

Now that we have introduced our conceptual framework using a fictional example, this section will describe our pipeline for acquiring relative posture time series data from video recording. To represent the movement of ferrets as a postural time series, we used DeepLabCut^10,12^, a markerless motion tracking software to track the pixel locations of animal body segments from video recording (Figure 2A). While the DeepLabCut produced time series demonstrates how posture changes over time, this representation shows only the pixel location rather than the inter-animal interaction. Instead of absolute location, we used relative posture metrics as input to Temporal Mapper. We quantified the interaction between both animals in three main ways: relative distance, velocity, and angle between body segments (Figure 2B). Additionally, this existing dataset includes four human-annotated behavioral categories for each frame: social, non-social, animal 1 active, and animal 2 active (Figure 2C). These human-annotated behavioral categories were needed to demonstrate the relationship between moment-to-moment relative posture dynamics and slower-switching macroscopic behavioral states within the transition networks. Although these four behavioral categories can be used to analyze behavior, we created more refined partitions, which separate social and non-social behavior based on activity level. Here, we introduced three activity levels: *active* (animal 1 and 2 active), *unilateral* (animal 1 or 2 active), *sleep* (animal 1 and 2 inactive). Therefore, the existing four manual codings turned into six behavioral categories: *active social*, *active non-social*, *unilateral social*, *unilateral non-social*, *social sleep*, and *non-social sleep* (Figure 2D).

**Figure 2:**
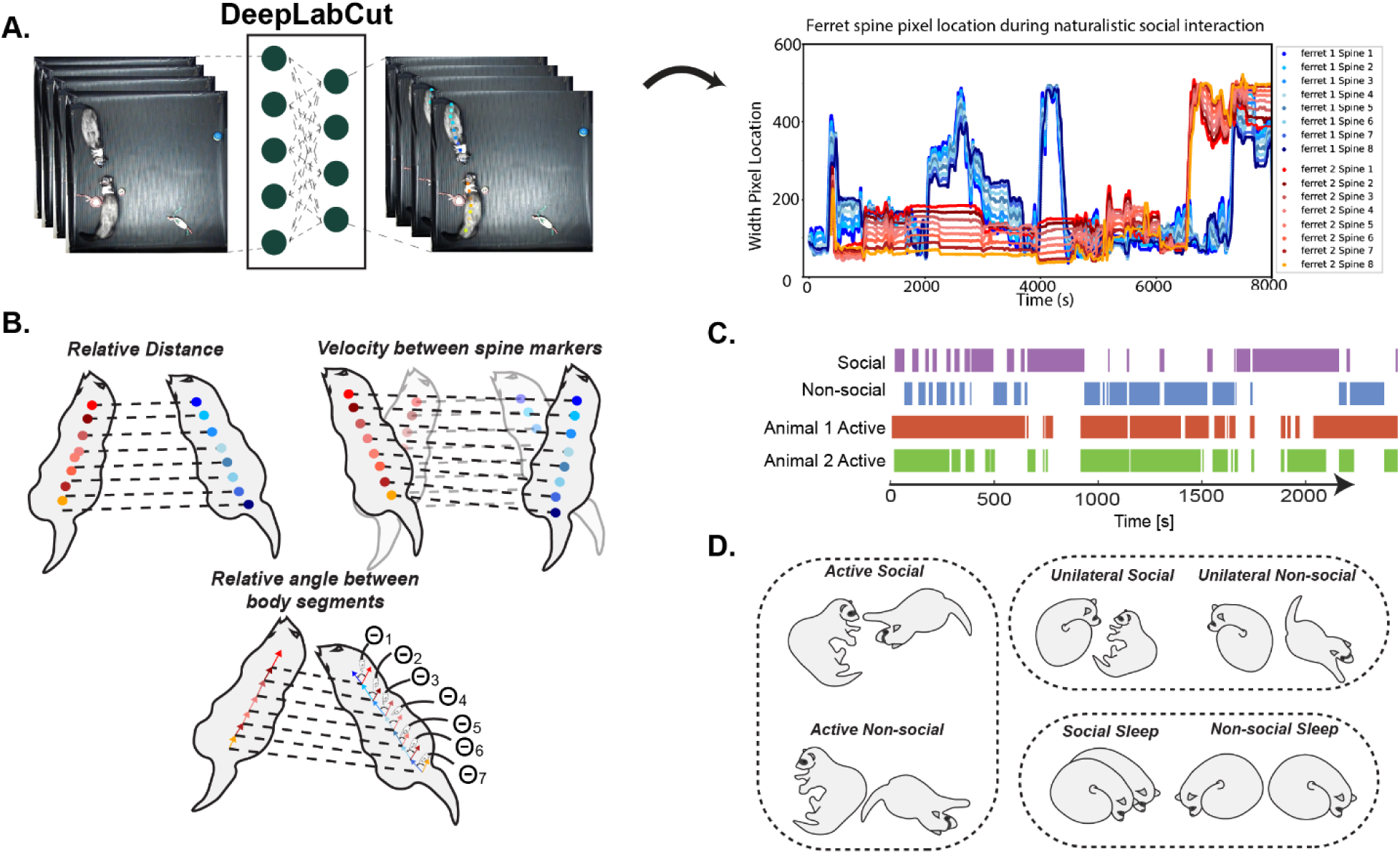
Relative posture time series for network construction and human annotated behavioral categories. (A) Posture dynamics were captured using DeepLabCut, a marker-less motion tracking software, to capture eight spine points in the bodies of two ferrets over time. Objects on the right side of the arena are toys. (B) Behavioral states were defined as relative postures based on three behavioral metrics: relative distance, relative velocity between spine markers, and relative angle between body segments. All three metrics were computed between homologous and non-homologous spine locations between two ferrets. (C) All behavioral sessions were human annotated at 30 Hz with four behavioral labels: social, non-social, animal 1 active, animal 2 active. (D) For the current work, social behavior was analyzed within the three activity levels: active, unilateral, and sleep. Labels were created automatically using the human annotated behavioral features in (C): *Active social* = (Animal 1 Active and Animal 2 Active) and Social. *Unilateral Social* = (Animal 1 Active or Animal 2 Active) and Social. *Social Sleep* = (Animal 1 not active and Animal 2 not active) and Social. Non-social equivalents are constructed with similar formula yet substituted “social” with “non-social.”

### Constructing and interpreting transition networks of relative postures

Using Temporal Mapper, a posture transition network was computed for each session from relative posture time series data (see one example in Figure 3A). The nodes represent distinct postural states (for relative posture metrics, see Figure 2B). The network edges are transitions between states. Node size indicates the behavioral state dwell time, where each node maps to a set of individual time points from the relative postural time series. The nodes were colored based on one of six behavioral categories described in Figure 2D. Nodes clustered together indicate close temporal connection, i.e., it takes fewer transitions for the animals to go from one posture state to another (see Methods for details). All discussed transition networks are high-dimensional objects where the 2-dimensional projections illustrated throughout this work are for visualization purposes only. An example path of connected nodes represents a sequence of relative posture state (Figure 3B).

**Figure 3:**
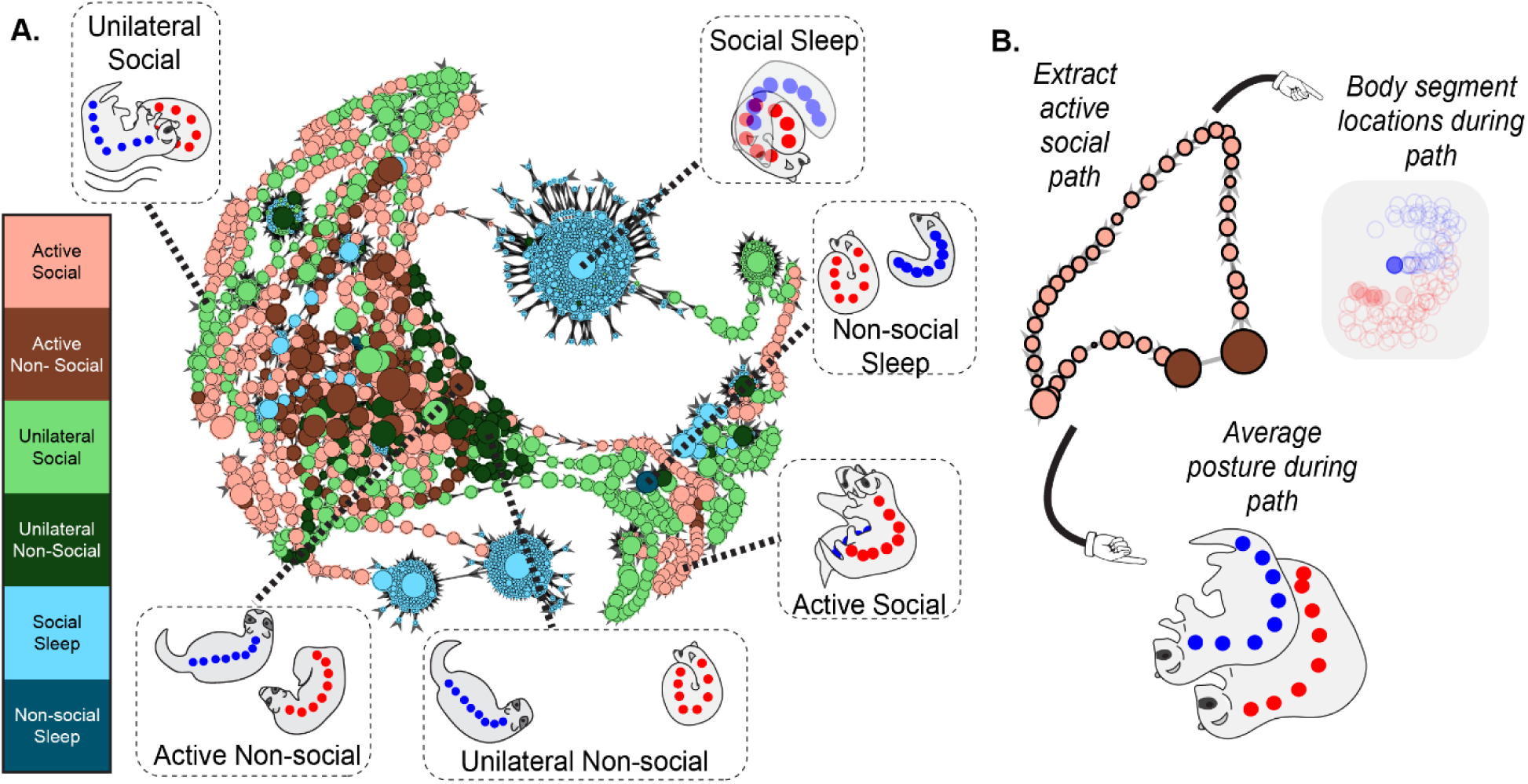
Interpreting the relative posture transition network. Transition networks capture the moment-to-moment behavioral dynamics represented as relative posture states (nodes) and their transitions (edges). The node colors reflect human-annotated behavioral categories. The node size represents the dwell time for each postural state. (A) Transition network from one example session with a duration of 35.75 minutes. Visually inspecting the periphery of the network, active social (salmon) and unilateral social (green) nodes are the dominant behavioral category present. Towards the center of the network are active non-social (dark salmon) and unilateral non-social (dark green) nodes. Social states are more widespread throughout the network, while non-social states exist closer to the center of the network. (B) Average posture of an active social path. Path extracted is an active social path where posture within each node can be averaged to form the average posture exhibited in the path.

### Temporal Mapper Reveals Widely Distributed and Brief Social States Throughout Transition Network

To further visualize the distribution of social nodes for direct comparisons, in Figure 4A1, we highlight the nodes by each behavioral category separately (Transition networks for all 12 sessions listed in Supplemental Figure 1). Visually, social nodes are distributed around the periphery of the network compared to the more centrally clustered non-social nodes. To quantify how widespread distribution of nodes is, we calculated the geodesic recurrence, which is a dissimilarity matrix consisting of pairwise distances (shortest path length) between states and depicts “how far away” states are from each other (Figure 4A2). We found that, on average, active social (** p<0.01; Figure 4B), unilateral social (*** p<0.001; Figure 4B), and social sleep (* p<0.05; Figure 4B) states are more widely distributed throughout the transition network compared to states within active non-social, unilateral non-social, and social non-sleep categories, respectively. This result suggests that social animals exhibit a wide range of postures during social interaction. Additionally, we found that active social nodes are significantly smaller compared to active non-social nodes (* p<0.05; Figure 4C). This observation suggests that active animals spend less time dwelling in the same posture during social compared to non-social activities. In short, these transition networks reveal that animals exhibit widespread, brief social states compared to centralized, longer-lasting non-social states.

**Figure 4:**
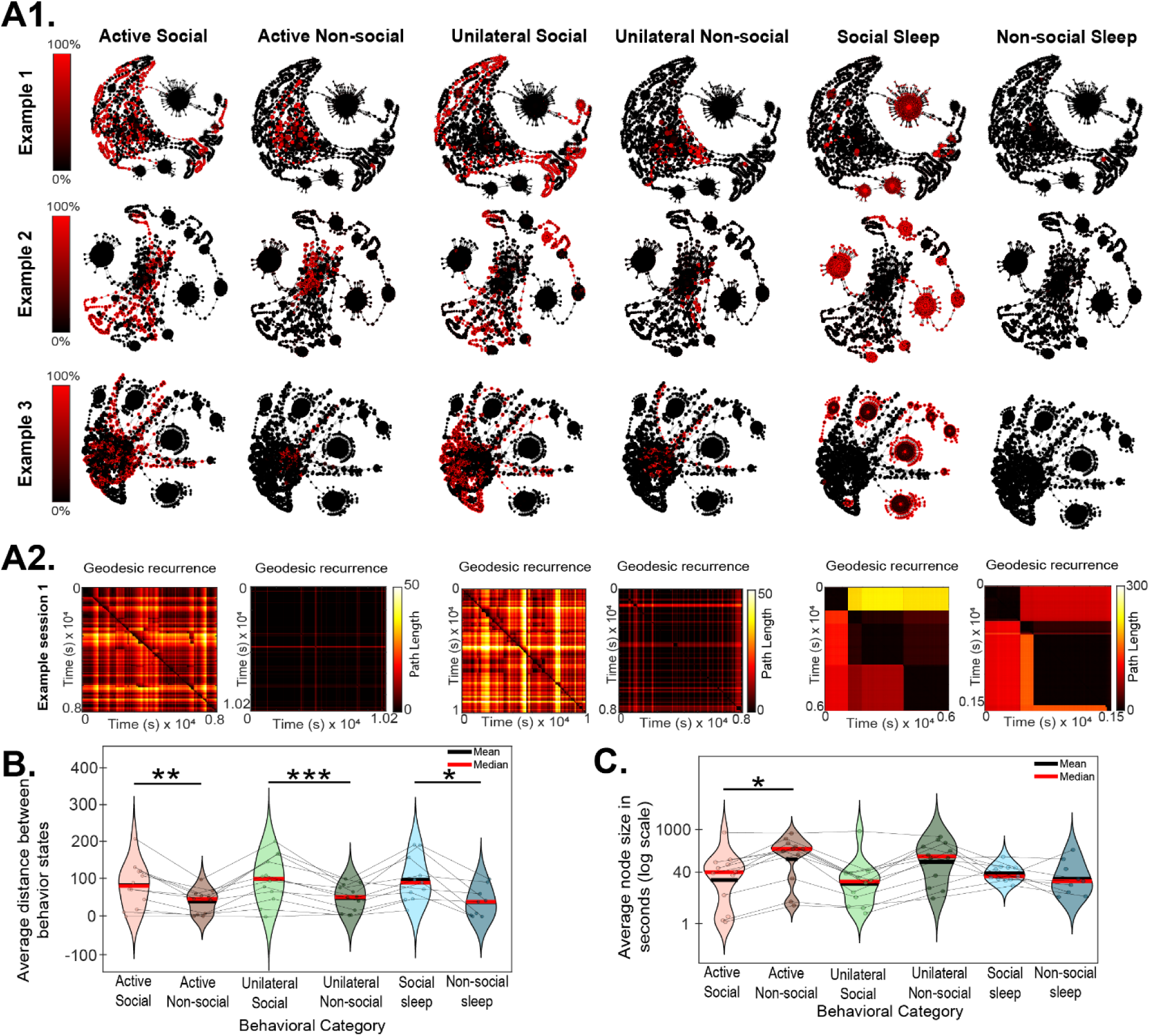
Social states are briefer and widely distributed throughout transition networks than non-social states. (A1) Three typical examples of a transition network for visual demonstration. Networks in each row have identical construction. The nodes are colored to highlight one of the six behavioral categories designated for each column. By visual inspection, social nodes are more widespread compared to non-social nodes. (A2) To quantify the widespread distribution in A1, Geodesic recurrence is computed, which is a dissimilarity matrix consisting of the shortest path length from state occupied at one time point to that of every other time point. Example Geodesic recurrence plots displayed for each behavioral category. Visually, social states are separated by longer paths compared to non-social states. (B) Across 12 sessions (average 26.96 minutes per session), social states are farther apart than non-social states across categories. (C) Active social nodes are significantly smaller than active non-social nodes across 12 sessions. Meaning, active animals spend significantly less time within social states compared to non-social states. (* p<0.05, ** p<0.01, *** p<0.001)

### Temporal Mapper Reveals Consistently Brief Posture State Sequences for Social Behavior within Transition Networks

In our analysis so far, we have examined the node distribution and size throughout transition networks for each recording session. This section focuses on behavioral state sequences, represented as paths in the transition network, which characterizes how relative posture states form sequences in social interaction. To examine the path construction unique to each behavior category in our transition network, we extracted sub-networks for each behavioral label where all the nodes were classified into the same behavioral category. Sub-networks for three example recording sessions shown in Figure 5A. Each behavioral category has its own unique collection of sub-networks where states are only connected to each other briefly in time. Our analysis demonstrates that social labeled sub-networks have more groups of paths, i.e., connected components, than non-social behaviors (Figure 5B). This result indicates that social states are more likely to form disconnected paths throughout the recording session. Visually, this conclusion matches the examples sub-networks illustrated (Figure 5A, sub-networks for all 12 sessions located in Supplementary Figure 2). Within active social sub-networks, there are many separate components whereas active non-social sub-networks have fewer components and are more connected. Additionally, two sample F-tests indicate that the spread of unilateral non-social path lengths is greater than unilateral social (Figure 5C). This result suggests that unilateral social transition networks have significantly more consistent path lengths across sessions (i.e., narrower distribution) compared to non-social transition network path lengths. Overall, these results suggest that animals in active social interaction performed a brief sequence of postures before transitioning to a different behavior. In particular, unilaterally social animals display a consistent number of sequential postures before changing behaviors.

**Figure 5:**
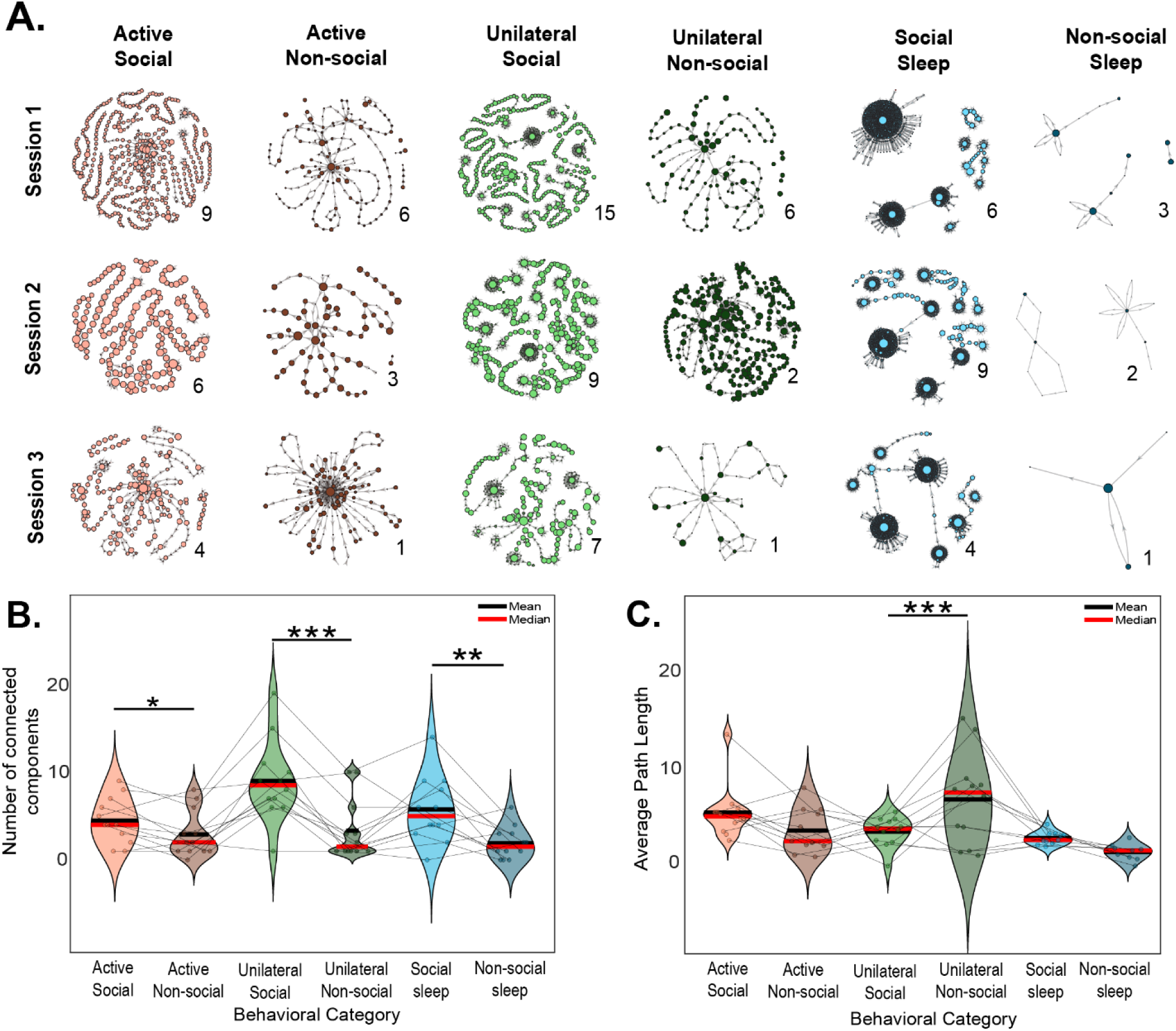
Social sub-networks are more disconnected than non-social sub-networks. (A) Behavioral sub-networks computed from three example sessions. Sub-networks from each behavioral category for illustration (same network used in Figure 2 and Figure 3). (B) Number of connected components labeled for each sub-network. Across all behavioral sub-networks in all 12 sessions, sub-networks with social behaviors have more connected components than sub-networks with non-social behaviors (* p<0.05, ** p<0.01, *** p<0.001, with paired-sample t-test). (C). Sub-networks for social behavior exhibit a significantly narrower distribution of path lengths, meaning unilaterally social animals perform a consistent number of postures before transitioning to a different behavior (***p<0.001, with a two-sample F-test).

## DISCUSSION

In the present work, we introduce Temporal Mapper as a novel tool for capturing rich behavioral dynamics during naturalistic social interaction. The effectiveness of Temporal Mapper for capturing complex, naturalistic social dynamics was demonstrated by application to characterizing ferret social interaction. In particular, we found that social states were widespread throughout postural transition networks compared to non-social states. In addition, we found that unilateral social states form sequences of more consistent length than their non-social counterparts.

### A comparison between related methods of social interaction and Temporal Mapper

In this study, we found that social states are brief and widespread throughout transition networks compared to non-social behavior. Our results are in alignment with existing findings that social animals display a wide range of social behaviors^9,46,47^. Previous studies standardized the classification of postures and manually documented up to 45 unique social postures across rat, mouse, hamster, and Guinea-pig^46^. Recent work extends standardized social postures by creating new methods that automatically classify behavior based on animal posture^15–24^ with specific work examining postural states and their transitions^16,33^. Specifically, MiceProfiler^16^ is a computational method to model and track mouse behavioral states and their transitions. In this work, de Chaumont and colleagues (2012) examined behavioral states and their transitions between socially interacting mice lacking neuronal nicotinic receptors compared to healthy C57BL/6J mice. Here, investigators found that the transition network constructed from healthy mice behaviors had more social states (8 social states) compared to non-social states (2 non-social states)^16^. In the current study, we did not find a significant difference between the number social versus non-social states.

However, this difference may be due to the limited number of states in Chaumont and colleagues’ transition network (10 states total) compared to the thousands of states utilized in the current study. Additionally, methods utilizing HMMs provide more coarse-grained representations of social behavior, with only one^22^ or two^22,33^ social states. By utilizing thousands of postural states, our current work was able to examine topological features of network construction, including the distance between nodes, their stability, and sub-networks, which was not investigated in social behavioral studies using HMMs^22,33^ or manually computed temporal graphs^16^. In our current work, Temporal Mapper revealed that social activity is arranged in consistent sequences of unstable postural states, which was previously not found in existing studies. Therefore, the current method provides a more network structure analysis of the behavioral states and their transitions in comparison to similar work.

### Potential Applications

The flexibility of Temporal Mapper makes our method well suited for characterizing rich dynamics in different species, scales, and paradigms. Temporal Mapper can be applied to various naturalistic recording sessions in indoor or outdoor settings and capture behavioral dynamics between single or multiple interacting animals. The high-dimensional transition networks from Temporal Mapper characterize the stability, similarity, and temporal linkage between behavioral states, which provides a new tool for modeling behavior with fine-grained characterization of complex, naturalistic dynamics. Beyond behavior, Temporal Mapper is applicable for capturing and connecting the moment-to-moment changes within neural dynamics. The flexible and data-driven design of Temporal Mapper allows for the identification of stable or unstable states and their transitions from neural time series data, such fMRI, EEG, and electrophysiology. In addition to animal settings, Temporal Mapper can be applied to human studies with investigations ranging from conflict management to patient-therapist interaction during psychotherapy settings. The versatility of Temporal Mapper provides plethora of applications for which our method can be applied to complex, dynamic naturalistic data.

### Limitations

The presented work comes with limitations. First, there is no established quantitative procedure for determining the parameters used in transition network construction.

Instead, tuning parameters are determined subjectively, better suited for exploratory rather than confirmatory analysis. Second, due to the complexity of this tool, the interpretation of nodes and edges is more complex. Traditional tools reduce the dimensionality of social behavior, which provides a simpler explanation for observed behavior. Since Temporal Mapper utilizes high-dimensional input for social behavior, the interpretation is inherently more complex and does not yield a simplistic explanation for social interaction. More exploratory analyses need to be completed to find converging interpretations and methods to derive generalizable, behaviorally relevant network features.

## Conclusion

In conclusion, Temporal Mapper captures rich behavioral dynamics observed during naturalistic social interaction. Complementing existing studies modeling behavioral dynamics, our present work provides an approach to capture the moment-to-moment behavioral states and their changes in naturalistic settings. This unique approach allows for the characterization of the global topological organization of postural states during social interaction, providing a direct link between postural dynamics and observed behavior. Additionally, Temporal Mapper has been utilized within task-based human neural analysis^34^ and real-world social dynamics^43^, which provides a common framework for analyzing complex dynamics within the brain and behavior. Future work will compare and bridge the behavioral and neural activity during naturalistic social interaction which captures the brain-to-behavior relationship between states and their transitions.

## MATERIALS AND METHODS

### Dataset used for method demonstration

#### Overview of the dataset

In this work, we analyzed an existing dataset of animal naturalistic social interaction between two domestic ferrets (*[Mustela putorius furo](https://www.sciencedirect.com/unsupported_browser)*). This dataset contains 12 sessions (26.96 ± 13.67 minutes) of video and electrophysiology recording. Additionally, four behavior codings were manually annotated at 10 Hz: social, non-social, animal 1 active, animal 2 active. The present demonstration only utilizes the behavioral data; analysis of the electrophysiological recordings will be the subject of a separate study.

#### Animals

Two spayed female ferrets were purchased from Marshal BioResources, North Rose, NY at 16-19 weeks old and weighing 0.7-1 kg. Both ferrets were grouped housed in 12 hours light/dark cycle. Procedures were performed in compliance with the Institutional Animal Care and Use Committee at the University of North Carolina at Chapel Hill and the United States Department of Agriculture (USDA Animal Welfare).

#### Video recording of free social interaction

Animals participated in naturalistic social interaction sessions where animals moved freely in a 3 ft x 3 ft behavioral arena. Toys for the animals were placed in arena as an option for non-social behavior. 12 sessions were recorded with an average session duration of 26.96 ± 13.67 minutes. Video recording was captured with a standard webcam (Spedal 920PRO Live Camera Full HD) with 1920x1080 resolution at 60 fps, mounted on the top center of the arena, pointing directly down. Top-down video footage was used to track body segment location via computer vision. Videos were downsampled to 30 fps, decreased resolution to 420p, and cropped to 540x540 pixels.

### Extracting Relative Postural Time Series from Video Recording

#### Behavioral Motion Tracking

Ferret spine positions during naturalistic social interaction were captured with DeepLabCut^10,12^, an open-source marker-less motion tracking software to record body movement from videos. We trained the DeepLabCut convolutional neural network to track the movement of 8 spine locations along both ferrets. We extracted ∼20 frames from each video session and manually annotated 8 spine locations on both animals. A total of 240 frames were used to train DeepLabCut to track ferret spine movement. The DeepLabCut ResNet-50 based neural network with default parameters and maximum iteration of 100,000 steps was trained on a workstation with intel i9 24-core CPU and NVIDIA GeForce RTX 4070 Ti GPU.

#### Behavioral Time Series Cleaning

The output from DeepLabCut provided the position of each spine marker of each animals within the behavioral chamber. DeepLabCut generated a decoded time series and subsequently low band-pass filtered (Python scipy.signal, 4th order Butterworth filter, 0.11 Hz cutoff). Automatically labeled frames were visually inspected, and labels were corrected as necessary. Manual correction was most needed when animals climbed on each other and occluded body segments. Semi-automatic corrections were applied to 5.769% ± 1.764 of frames.

#### Behavioral coding and social comparison categories

Behavioral sessions were coded with human annotated behavioral variables evaluated at 10 Hz throughout all sessions. The four codings included social, non-social, animal 1 active, animal 2 active. Animals were considered social based on the judgement of the human annotator. To be considered active, the animal must be moving within the arena. The analysis in this study focused on the social behavior between animals. Nevertheless, postural dynamics were also influenced by activity level. Therefore, comparisons between social vs non-social behaviors were only made at matched activity levels. Three dyadic activity levels are derived from human-annotated variables: (1) *active* where both animals are active, (2) *unilateral* active where only one animal is active, and (3) *sleep* where both animals are inactive.

#### Relative posture time series for network construction

To quantify the interpersonal interaction between ferrets, relative postures were computed from absolute spine position time series. The relative posture was defined in terms of the relative distance, relative velocity between spine markers, and relative angle between body segments. Relative distance measured the pairwise Euclidean distance between spine markers. Relative velocity was calculated by subtracting positions from the current time step from the previous time step and divided by the time step duration (1/30 seconds). Relative angles were calculated between the body segments of the animals. All metrics were calculated using homologous and non-homologous spine markers between animals.

### Constructing relative postural transition networks using Temporal Mapper

*Temporal Mapper network construction.* Computing transition networks with Temporal Mapper requires 3 steps: 1) compute pairwise distance matrix ***D*** from time series data ***X****[t]*, where the rows in ***X*** correspond to time steps and columns correspond to state variables (relative posture variable in the present case); 2) Compute a spatiotemporal k-nearest neighbor (stknn) graph from the pairwise distance matrix ***D*** and the temporal order of sample points. The stknn graph is a transition network where nodes correspond to time points within the time series data and the edges are the transitions between states. Nodes connected with a single edge are sequential in time. Bidirectionally connected nodes are near each other in the state space, where *k* is the spatial neighborhood for each node. In the current work, we used *k* = 10 as the spatial neighborhood for each transition network. 3) Compute a simplified network from stknn by compressing nodes connect to each other within distance *d* in the stknn to a single node in the simplified network, where distance *d* is measures as the shortest path length between nodes in the stknn. Here, we used *d* = 3 to construct all simplified postural transition networks. Networks nodes were colored based on the average expressed behavioral attribute (active social, active non-social, unilateral social, unilateral non-social, social sleep, or non-social sleep; Figures 3-5) during the time steps within the corresponding node. All long behavioral sessions were capped at 38.88 minutes (70,000 time-steps) due to memory constraints.

#### Transition network visualization

All computations and statistical measures with respect to transition networks were completed in the original, high-dimensional space. 2-dimensional projects of transition networks within Figures 3-5 and Supplemental Figures 1-2 are for visual illustration only. For visualization, transition networks are plotted in MATLAB using force-directed placement^48^.

#### Geodesic recurrence plot

Geodesic recurrence plots ***G*** capture the shortest path between nodes within a directed network in a time-resolved manner. A geodesic recurrence plot is N-by-N dissimilarity matrix, where N is the number of time points and the element ***G****i,j* is the minimal path length between nodes occupied at any two time points *i* and *j* within the transition network. The path lengths are generally not symmetric since the transition networks are directed. The Geodesic recurrence within each behavioral category was calculated with the following equation: 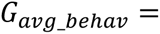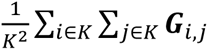 where *K* is the set of all time points for a behavior category.

#### Sub-network extraction

First, we compute the average expressed behavioral category expressed for each node in the transition network. Second, we exacted subgraphs of the full transition networks, resulting in new transition networks where all nodes correspond to the same behavioral category. Edges only connect nodes that are connected in time.

### Statistical analysis

Paired t-tests were used to determine the statistical significance of average node size between social and non-social categories. Additionally, paired t-tests were used to determine the statistical significance between average node distances within the transition network between social and non-social categories. Two sample F-tests were used to determine whether variances in path lengths between social and non-social were equal in each activity level.

## AUTHOR CONTRIBUTIONS

Jared Reiling: Formal analysis; Investigation; Methodology; Software; Visualization; Writing—original draft; Writing—review & editing. Nancy Padilla-Coreano: Writing—review & editing. Dhruvi Patel: Data collection; Data analysis—behavior coding. Flavio Frohlich: Funding acquisition; Writing—review & editing. Mengsen Zhang: Conceptualization; Funding acquisition; Investigation; Methodology; Project administration; Supervision; Validation; Writing—review & editing.

---

**Supplemental Figure 1:**
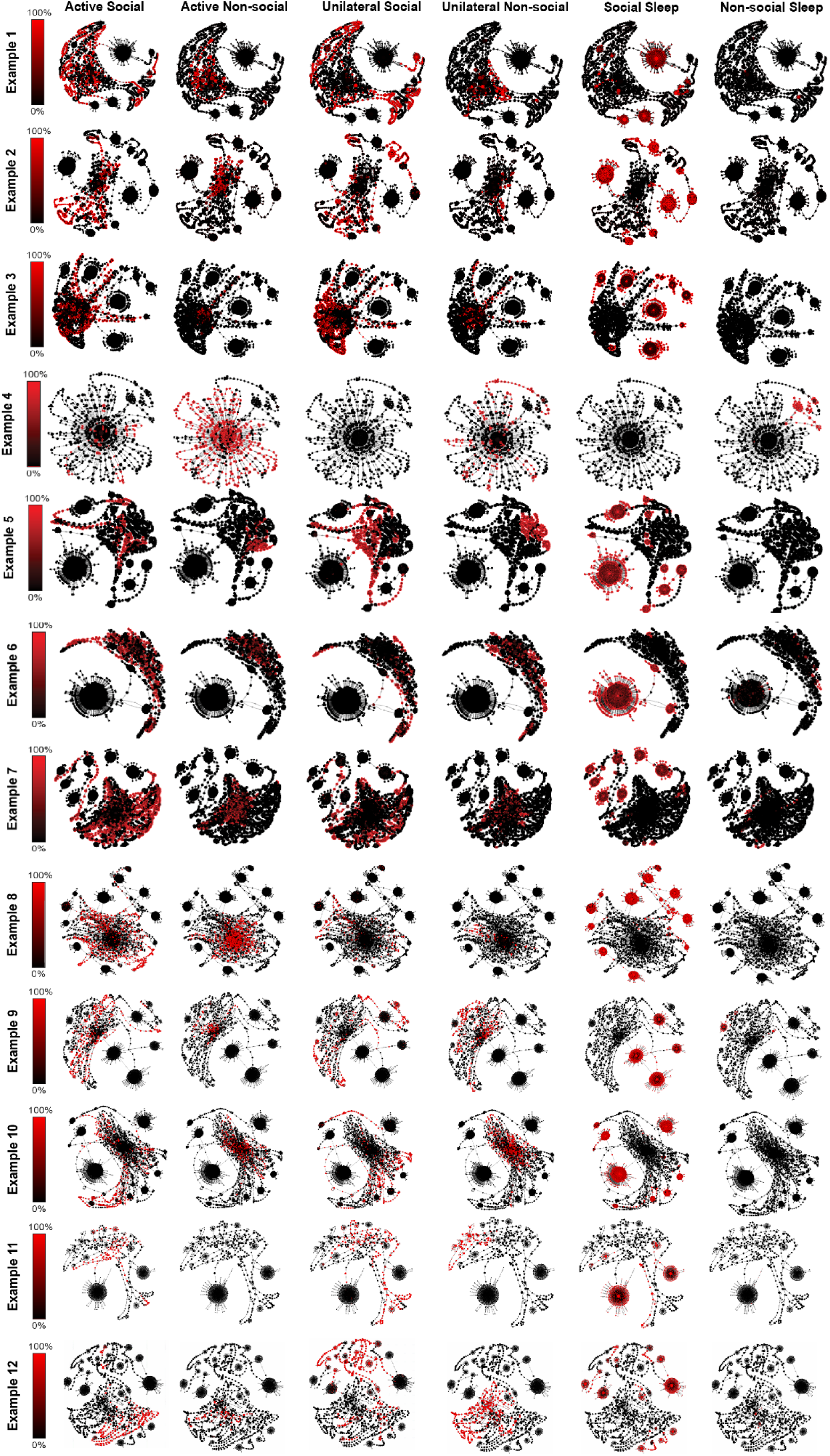
Transition networks of each behavioral session. 12 behavioral sessions analyzed. The networks in each row have identical construction and color based on the indicated behavioral category for each column.

**Supplemental Figure 2:**
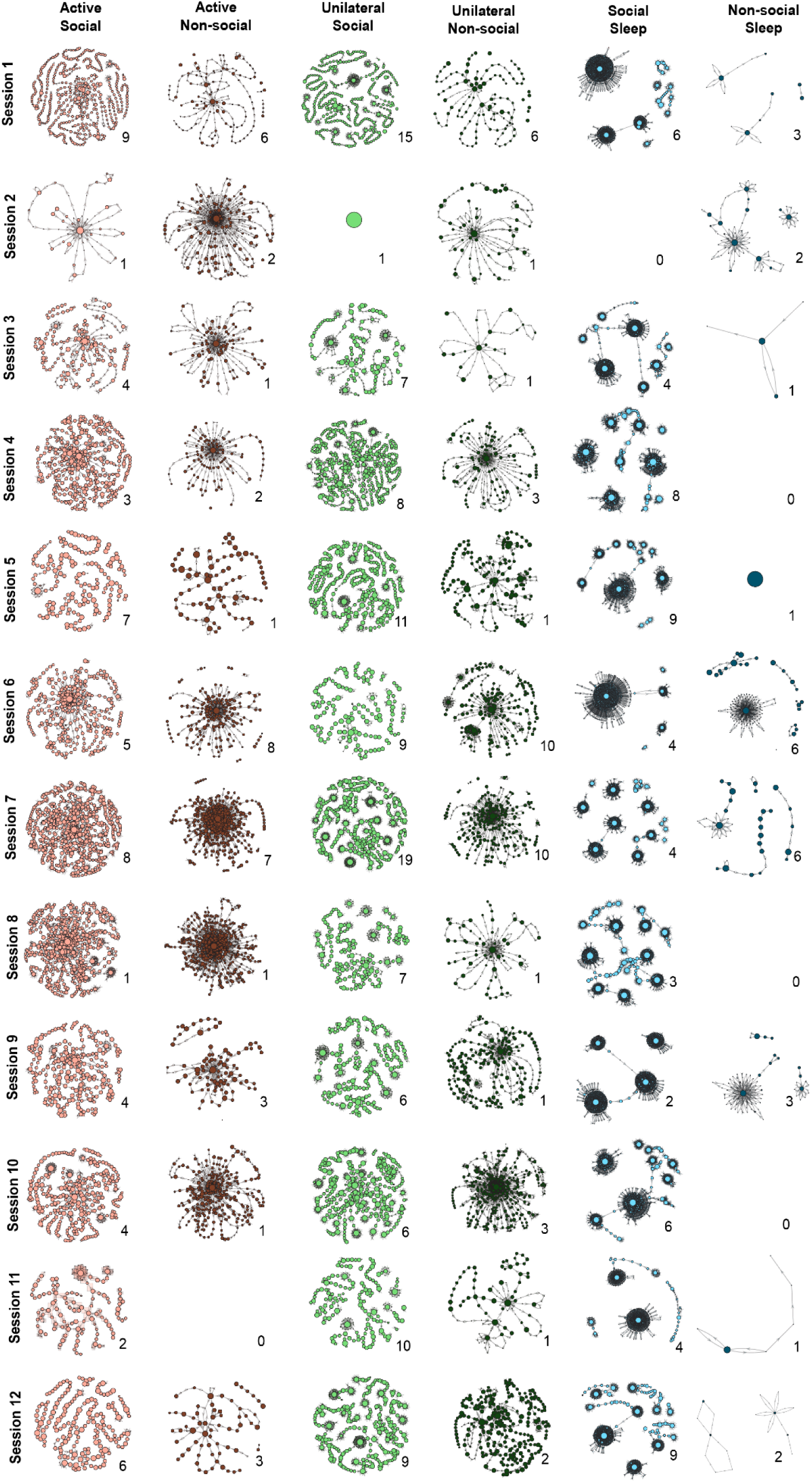
Behavioral sub-networks computed from all 12 analyzed sessions. Number next to each sub-network indicates the number of connected components.

## REFERENCES

1. Miller, J. H. & Page, S. E. Complex Adaptive Systems: An Introduction to Computational Models of Social Life. (Princeton University Press, 2007).

2. Bolis, D., Dumas, G. & Schilbach, L. Interpersonal attunement in social interactions: from *collective* psychophysiology to *inter- personalized* psychiatry and beyond. Philos. Trans. R. Soc. B Biol. Sci. 378, 20210365 (2023).

3. Dumas, G., Lachat, F., Martinerie, J., Nadel, J. & George, N. From social behaviour to brain synchronization: Review and perspectives in hyperscanning. IRBM 32, 48–53 (2011).

4. Moy, S. S. et al. Sociability and preference for social novelty in five inbred strains: an approach to assess autistic-like behavior in mice. Genes Brain Behav. 3, 287–302 (2004).

5. Oliva, A., Fernández-Ruiz, A., Leroy, F. & Siegelbaum, S. A. Hippocampal CA2 sharp-wave ripples reactivate and promote social memory. Nature 587, 264–269 (2020).

6. Gheusi, G., Bluthé, R.-M., Goodall, G. & Dantzer, R. Social and individual recognition in rodents: Methodological aspects and neurobiological bases. Behav. Processes 33, 59–87 (1994).

7. Krueger-Burg, D. et al. The SocioBox: A Novel Paradigm to Assess Complex Social Recognition in Male Mice. Front. Behav. Neurosci. 10, (2016).

8. De Paula Cunha Almeida, C., et al. Prefrontal and Subcortical c-Fos Mapping of Reward Responses across Competitive and Social Contexts. eneuro 12, ENEURO.0158-25.2025 (2025).

9. File, S. E. & Hyde, J. R. Can social interaction be used to measure anxiety? Br. J. Pharmacol. 62, 19–24 (1978).

10. Mathis, A. et al. DeepLabCut: markerless pose estimation of user-defined body parts with deep learning. Nat. Neurosci. 21, 1281–1289 (2018).

11. Nath, T. et al. Using DeepLabCut for 3D markerless pose estimation across species and behaviors. Nat. Protoc. 14, 2152–2176 (2019).

12. Lauer, J. et al. Multi-animal pose estimation, identification and tracking with DeepLabCut. Nat. Methods 19, 496–504 (2022).

13. Goodwin, N. L. et al. Simple Behavioral Analysis (SimBA) as a platform for explainable machine learning in behavioral neuroscience. Nat. Neurosci. 27, 1411–1424 (2024).

14. Karashchuk, P. et al. Anipose: A toolkit for robust markerless 3D pose estimation. Cell Rep. 36, 109730 (2021).

15. Goodwin, N. L. et al. Simple Behavioral Analysis (SimBA) as a platform for explainable machine learning in behavioral neuroscience. Nat. Neurosci. 27, 1411–1424 (2024).

16. de Chaumont, F. et al. Computerized video analysis of social interactions in mice. Nat. Methods 9, 410–417 (2012).

17. Kabra, M., Robie, A. A., Rivera-Alba, M., Branson, S. & Branson, K. JAABA: interactive machine learning for automatic annotation of animal behavior. Nat. Methods 10, 64–67 (2013).

18. Giancardo, L. et al. Automatic Visual Tracking and Social Behaviour Analysis with Multiple Mice. PLOS ONE 8, e74557 (2013).

19. Segalin, C. et al. The Mouse Action Recognition System (MARS) software pipeline for automated analysis of social behaviors in mice. eLife 10, e63720 (2021).

20. Bohnslav, J. P. et al. DeepEthogram, a machine learning pipeline for supervised behavior classification from raw pixels. eLife 10, e63377 (2021).

21. Harris, C., Finn, K. R., Kieseler, M.-L., Maechler, M. R. & Tse, P. U. DeepAction: a MATLAB toolbox for automated classification of animal behavior in video. Sci. Rep. 13, 2688 (2023).

22. Arakawa, T. et al. Automated Estimation of Mouse Social Behaviors Based on a Hidden Markov Model. in Hidden Markov Models: Methods and Protocols (eds Westhead, D. R. & Vijayabaskar, M. S.) 185–197 (Springer, New York, NY, 2017). doi:10.1007/978-1-4939-6753-7_14.

23. de Chaumont, F. et al. Real-time analysis of the behaviour of groups of mice via a depth-sensing camera and machine learning. *Nat*. Biomed. Eng. 3, 930–942 (2019).

24. Chen, Z. et al. AlphaTracker: a multi-animal tracking and behavioral analysis tool. Front. Behav. Neurosci. 17, (2023).

25. Tofani, G. S. S. et al. Gut microbiota regulates stress responsivity via the circadian system. Cell Metab. 37, 138–153.e5 (2025).

26. Bordes, J. et al. Automatically annotated motion tracking identifies a distinct social behavioral profile following chronic social defeat stress. Nat. Commun. 14, 4319 (2023).

27. Morales, J. M., Haydon, D. T., Frair, J., Holsinger, K. E. & Fryxell, J. M. Extracting More Out of Relocation Data: Building Movement Models as Mixtures of Random Walks. Ecology 85, 2436–2445 (2004).

28. Patterson, T. A., Basson, M., Bravington, M. V. & Gunn, J. S. Classifying movement behaviour in relation to environmental conditions using hidden Markov models. J. Anim. Ecol. 78, 1113–1123 (2009).

29. Langrock, R. et al. Flexible and practical modeling of animal telemetry data: hidden Markov models and extensions. Ecology 93, 2336–2342 (2012).

30. Langrock, R., Marques, T. A., Baird, R. W. & Thomas, L. Modeling the Diving Behavior of Whales: A Latent-Variable Approach with Feedback and Semi-Markovian Components. J. Agric. Biol. Environ. Stat. 19, 82–100 (2014).

31. Michelot, T., Langrock, R. & Patterson, T. A. moveHMM: an R package for the statistical modelling of animal movement data using hidden Markov models. Methods Ecol. Evol. 7, 1308–1315 (2016).

32. Leos-Barajas, V. et al. Multi-scale Modeling of Animal Movement and General Behavior Data Using Hidden Markov Models with Hierarchical Structures. J. Agric. Biol. Environ. Stat. 22, 232–248 (2017).

33. Stednitz, S. J. et al. Coordinated social interaction states revealed by probabilistic modeling of zebrafish behavior. Curr. Biol. 35, 2903–2915.e6 (2025).

34. Zhang, M., Chowdhury, S. & Saggar, M. Temporal Mapper: Transition networks in simulated and real neural dynamics. Netw. Neurosci. 7, 431–460 (2023).

35. Carlsson, G. Topology and Data. Bull. Am. Math. Soc. - BULL AMER MATH SOC 46, 255–308 (2009).

36. Munch, E. A User’s Guide to Topological Data Analysis. J. Learn. Anal. 4, 47–61 (2017).

37. Amézquita, E. J., Quigley, M. Y., Ophelders, T., Munch, E. & Chitwood, D. H. The shape of things to come: Topological data analysis and biology, from molecules to organisms. Dev. Dyn. 249, 816–833 (2020).

38. Saggar, M. et al. Towards a new approach to reveal dynamical organization of the brain using topological data analysis. Nat. Commun. 9, 1399 (2018).

39. Geniesse, C., Sporns, O., Petri, G. & Saggar, M. Generating dynamical neuroimaging spatiotemporal representations (DyNeuSR) using topological data analysis. Netw. Neurosci. 3, 763–778 (2019).

40. Zhang, M., Kalies, W. D., Kelso, J. A. S. & Tognoli, E. Topological portraits of multiscale coordination dynamics. J. Neurosci. Methods 339, 108672 (2020).

41. Curto, C. What can topology tell us about the neural code? Bull. Am. Math. Soc. 54, 63–78 (2016).

42. Singh, G., Mémoli, F. & Carlsson, G. Topological Methods for the Analysis of High Dimensional Data Sets and 3D Object Recognition. in 91–100 (2007). doi:10.2312/SPBG/SPBG07/091-100.

43. Luo, X. & Zhang, M. A topological data analysis method for revealing dynamic changes in psychotherapy microprocesses. Front. Psychol. 16, (2026).

44. Hernádi, A., Kis, A., Turcsán, B. & Topál, J. Man’s Underground Best Friend: Domestic Ferrets, Unlike the Wild Forms, Show Evidence of Dog-Like Social-Cognitive Skills. PLOS ONE 7, e43267 (2012).

45. Boyce, S. W., Zingg, B. M. & Lightfoot, T. L. Behavior of *Mustela Putorius Furo* (The Domestic Ferret). Veterinary Clin. North Am. Exot. Anim. Pract. 4, 697–712 (2001).

46. Grant, E. C. & Mackintosh, J. H. A Comparison of the Social Postures of Some Common Laboratory Rodents. Behaviour 21, 246–259 (1963).

47. Poole, T. B. & Fish, J. An investigation of playful behaviour in Rattus norvegicus and Mus musculus (Mammalia). J. Zool. 175, 61–71 (1975).

48. Fruchterman, T. M. J. & Reingold, E. M. Graph drawing by force-directed placement. Softw. Pract. Exp. 21, 1129–1164 (1991).

